# Hierarchical modeling of mechano-chemical dynamics of epithelial sheets across cells and tissue

**DOI:** 10.1101/859348

**Authors:** Yoshifumi Asakura, Yohei Kondo, Kazuhiro Aoki, Honda Naoki

## Abstract

Collective cell migration is a macroscopic population-level phenomenon that has emerged across hierarchy from microscopic interactions between cells; however, the underlying mechanism remains unclear. Here, we targeted epithelial collective cell migration, driven by the mechanical force regulated by chemical signals of traveling ERK activation waves, observed in wound healing. We propose a hierarchical mathematical framework for understanding how cells are orchestrated through mechanochemical cell-cell interaction. We mathematically transformed a particle-based model at the cellular level into a continuum model at the tissue level. The continuum model described relationships between cell migration and mechanochemical variables, namely, ERK activity gradients, cell density, and velocity field, which could be compared with live-cell imaging data. The continuum model recapitulated the ERK wave-induced collective cell migration in wound healing. We also numerically confirmed a consistency between the two models. This framework allows us to connect hierarchical causality from the single-cell level to the tissue level.

## Introduction

Collective cell migration in epithelial sheets plays an important role in embryonic development and tissue homeostasis [1]. Such a tissue-level phenomenon must have emerged through interactions between individual cells. However, how the intercellular interactions affect collective cell migration remains unclear, because of the difficulties in understanding causality between the microscopic intercellular process and the macroscopic tissue-level phenomenon. In this study, we aimed to address this issue through computational modeling across the hierarchy between cells and tissues.

Several studies have investigated the molecular basis of chemical and mechanical regulation of cell migration. Recently, it was revealed that actomyosin-generated mechanical force is regulated by chemical cues such as E-cadherin [2] and Rho GTPases [3, 4] during collective cell migration. In addition, extracellular signal-related kinase (ERK), involved in the MAPK cascade plays an important role in single cell migration *in vitro* [5] and *in vivo* [6, 7], through phosphorylation of myosin light chain (MLC) kinase and focal adhesion kinase.

Recently, we reported that collective cell migration during wound healing was mediated by mechano-chemical cellular dynamics involving ERK activation *in vivo* and *in vitro* [8, 9]. In these previous studies, we experimentally discovered that ERK activation spread in space and time as traveling waves during wound healing, as revealed by live-imaging of ERK activity using a biosensor based on the principle of fluorescence resonance energy transfer (FRET) [6, 7, 10]. ERK chemical waves spread from the site of injury across the epithelial sheet, and cells collectively move toward the site of injury; this implies that cell migration occurs in a direction opposite to that of ERK waves (Fig. 1). We also showed that cells altered their mechanical properties, including mobility and volume, in response to ERK activation.

**Figure 1:**
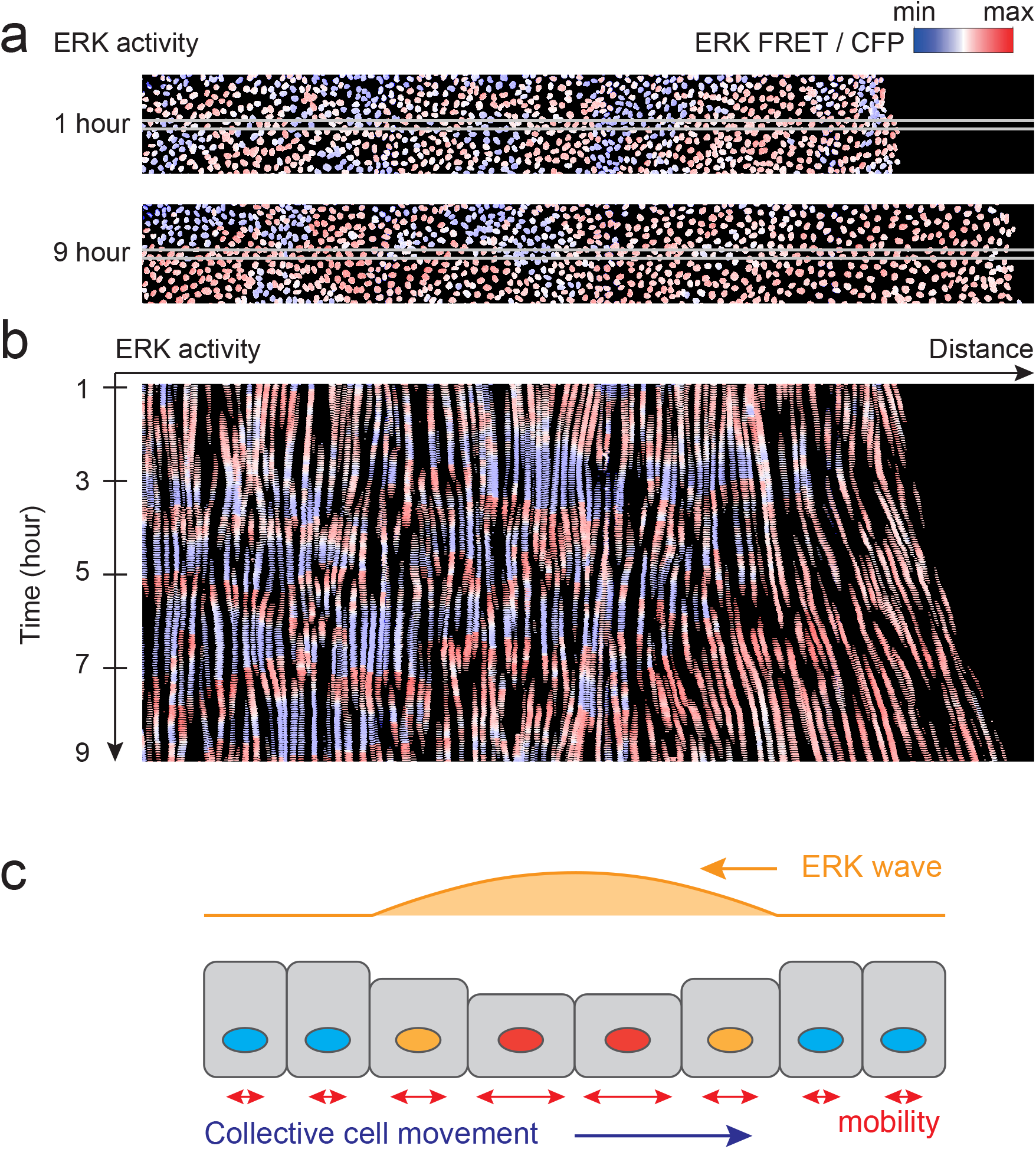
Live imaging of ERK-induced collective cell migration a. Visualization of ERK activity during wound-healing in epithelial MDCK cell sheet express-ing the FRET-based biosensor of ERK. Snapshots are presented at 1 hr (upper panel) and 9 hr (lower panel) after scratching. Red and blue colors indicate high and low ERK activity, respectively. b. A kymograph of ERK activity images in a band region of interest indicated by the white box in (a). c. A schematic representation of ERK-mediated collective cell migration. Cells migrate to-ward the opposite direction of the ERK wave. The cellular volume and mobility are in-creased by ERK activation.

To understand how the ERK activation wave is transformed to collective cell migration, we previously developed a particle-based model of an epithelial cell population [8]. This model was formulated as one-dimensionally aligned particles connected by springs, in which particles and springs represent cells and elastic interactions, respectively. According to our experiments, we incorporated the effects of ERK signals on cellular mechanical properties, namely ERK-dependent mobility and volume, in the model. Using computer simulations, our model successfully recapitulated the basic property of ERK-mediated collective cell migration, i.e., cells migrate in a direction opposite to that of the ERK traveling wave.

However, the particle-based model could not facilitate a direct comparison of the experimental data with the model, because of the following discrepancies. First, our previous model was implemented in one dimension whereas the epithelial tissue migrated in two dimensions. Second, even in a possible 2D model, it is hard to compare the model with cellular behaviors observed in live-imaging, owing to the plastic nature of the tissue. In the tissue, cellular positions are not stably fixed, as is observed in solid particles; it behaves like a fluid in which cellular positions are rearranged in time. Additionally, cells stochastically pass through other cells. Third, cellular responses are heterogeneous; however, our previous model assumed that cells are homogeneous with the same parameter values.

To overcome the aforementioned limitations, we aimed to acquire a coarse-grained model of the two-dimensional epithelial dynamics that averages the heterogeneous properties of individual cells. We propose a new hierarchical approach, in which our particle-based model of ERK-mediated collective cell migration was approximately translated into a continuum model. This approach allows us to seamlessly connect hierarchical causality from single-cell behavior to tissue-level dynamics. In the following sections, we describe the particle-based model and its transformation to the continuum model and demonstrate the validity of our hierarchical approach by comparing these two models via numerical simulations.

## Results

### Particle-based model with viscosity

Previously, we developed a particle-based model to understand how the ERK traveling wave mediates collective cell migration in epithelial sheets, in which cells adhere to neighboring cells so that they cannot freely migrate but rather mechanically interact each other through repulsive and attractive forces (Fig. 2a). In the model, the epithelial sheet was considered a one-dimensional (1D) series of particles connected through springs, where particles denote the positions of cells and springs represent the elastic property of the cells involved in the membrane, cytoskeleton and adhesion [8]. Thus, the dynamics of the cellular position *x*_*i*_ is described by the following ordinary differential equations (ODEs):

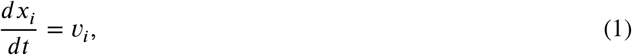

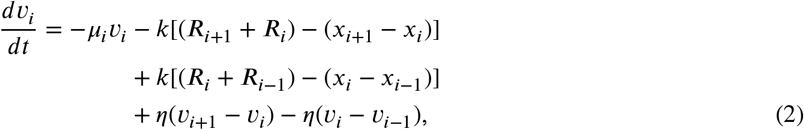

where *x*_*i*_ and *v*_*i*_ indicate the position and velocity of the i-th cell, respectively. Parameters *μ*_*i*_, *k*, *R*_*i*_, *η* indicate the friction coefficient, spring constant, cell radius, and viscosity coefficient, respectively. Because cell size and mobility were upregulated by ERK activation in our previous study [8], we assumed that cell radius and friction were positively or negatively modulated by ERK activity as

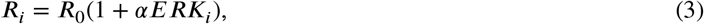

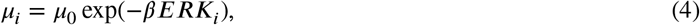

where *R*_0_ and *μ*_0_ indicate the basal radius and basal friction of the cells, respectively; *α* and *β* denote the effect of ERK activity on cellular size and friction, respectively. ERK activity was applied as traveling Gaussian distributions as

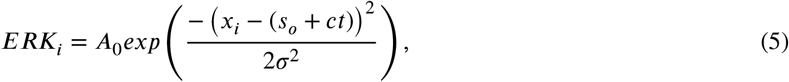

where *ERK*_*i*_, *A*_0_, *s*_*0*_, *c*, *σ*^2^ indicate the ERK activity of the *i*-th cell, amplitude of the ERK wave, initial position of an ERK wave, velocity of the ERK wave, and a positive constant regulating width of the ERK wave, respectively.

**Figure 2:**
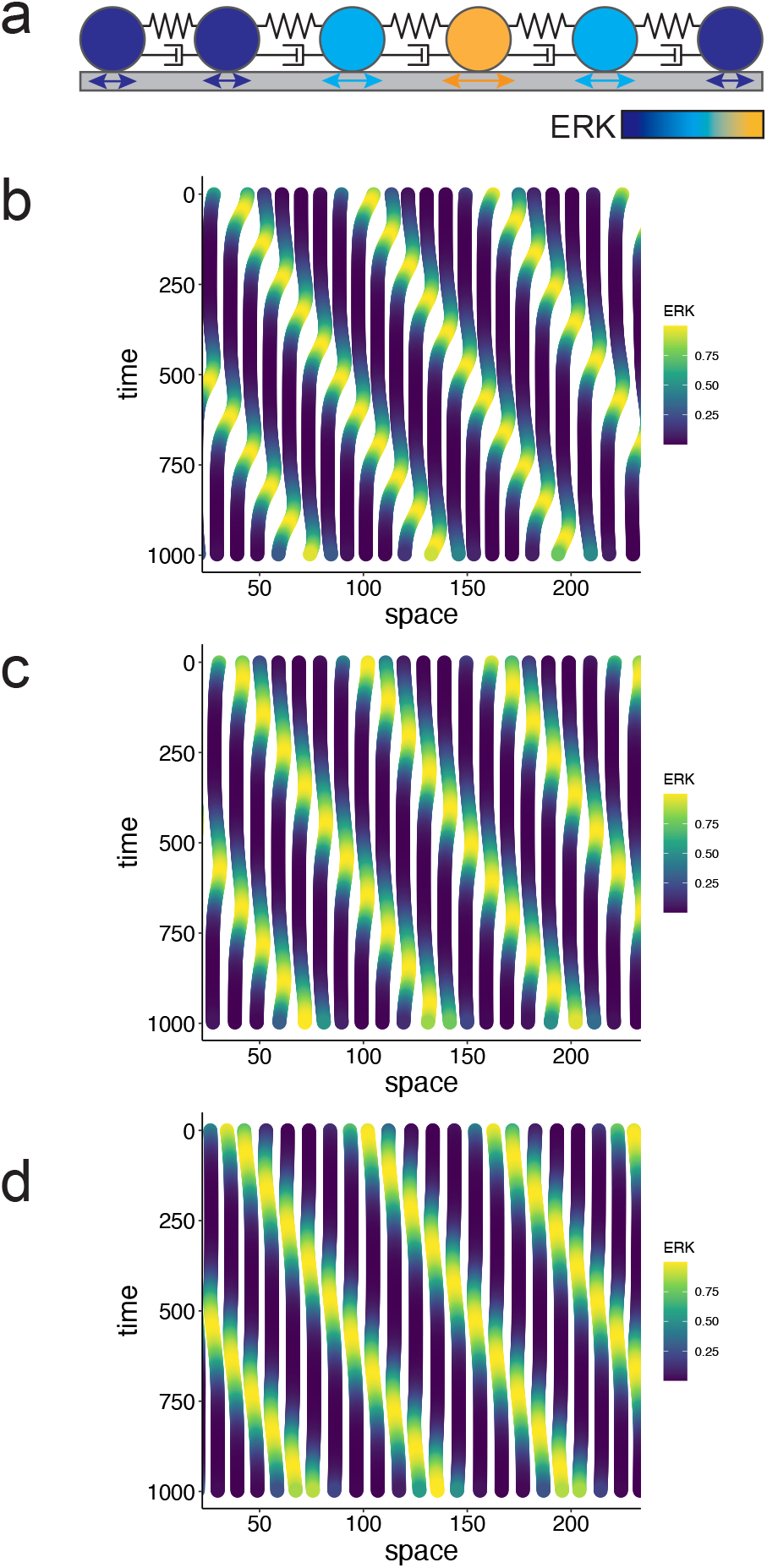
Simulations of the particle-based model in 1D a. Graphical representation of the particle-based model. The centers of cells are illustrated by particles, and these are connected with neighboring cells by springs and dampers representing the elasticity and viscosity of the cytoplasm, respectively. b-d. Simulations of ERK-induced collective cell migration with different parameter settings. Each line indicates cellular trajectory with the line color representing ERK activity. Parameters in Table 1 were used. In (b), cells migrate toward the opposite direction of the ERK waves. In (c), cells show back-and-forth movement during the ERK waves without net displacement. In (d), cells migrate toward the same direction of the ERK waves.

**Table 1:** List of parameter values

| variables | Fig2B,<br>Fig3C-E left,<br>Fig4A and Fig5 | Fig2C,<br>Fig3C-E middle,<br>Fig4B | Fig2D,<br>Fig3C-E right,<br>Fig4C |
| --- | --- | --- | --- |
| $\mu_0$ | 10 | 10 | 10 |
| $\alpha$ | 1.5 | 1.5 | -0.4 |
| $\beta$ | 2.5 | 0 | 2.5 |
| $k_0$ | 2 | 2 | 2 |
| $R_0$ | 0.5 | 0.5 | 0.5 |
| $\eta$ | 0.08 | 0.08 | 0.08 |
| $r$ | 1 | 1 | 1 |

Through the simulation, the modified model showed ERK-induced cell migration, in which the cells migrated in direction opposite to that of the ERK traveling wave (Fig. 2b); this was consistent with our previous experiments. dditionally, by changing the simulation parameters, migration in the opposite direction was abolished (Fig 2c, d). Cancelling the effects of ERK on friction coefficient *μ*_*i*_ abolished any migration after the passage of an ERK wave (Fig 2c), and the negative effect of ERK on cell radius *R*_*i*_ caused migration in the same direction as the ERK traveling wave (Fig 2d). These parameter dependencies of the direction of cellular migration validated our assumption of the effects of ERK on cellular properties expressed in equation (3) and (4).

### Continuum model

We coarse-grained the 1D particle-based model by continuum dynamics approximation (Fig 3a). We first transformed equation (2) into

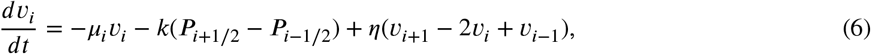

where *P*_*i*_ = (*R*_*i*+1/2_ + *R*_*i*−1/2_) − (*x*_*i*+1/2_ + *x*_*i*−1/2_). *P*_*i*−1/2_ and *P*_*i*+1/2_ correspond to the pressure from left and right neighboring cells. Since the cell density is defined as a number of cells within unit distance, the cell density at the middle position between the cells *i* and *i* ± 1 is calculated by

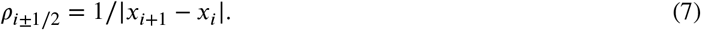

In the same way, natural density is defined by 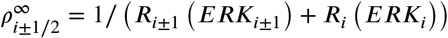 Thus,

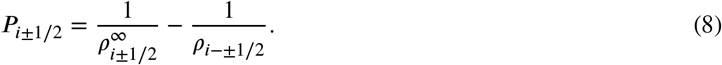

**Figure 3:**
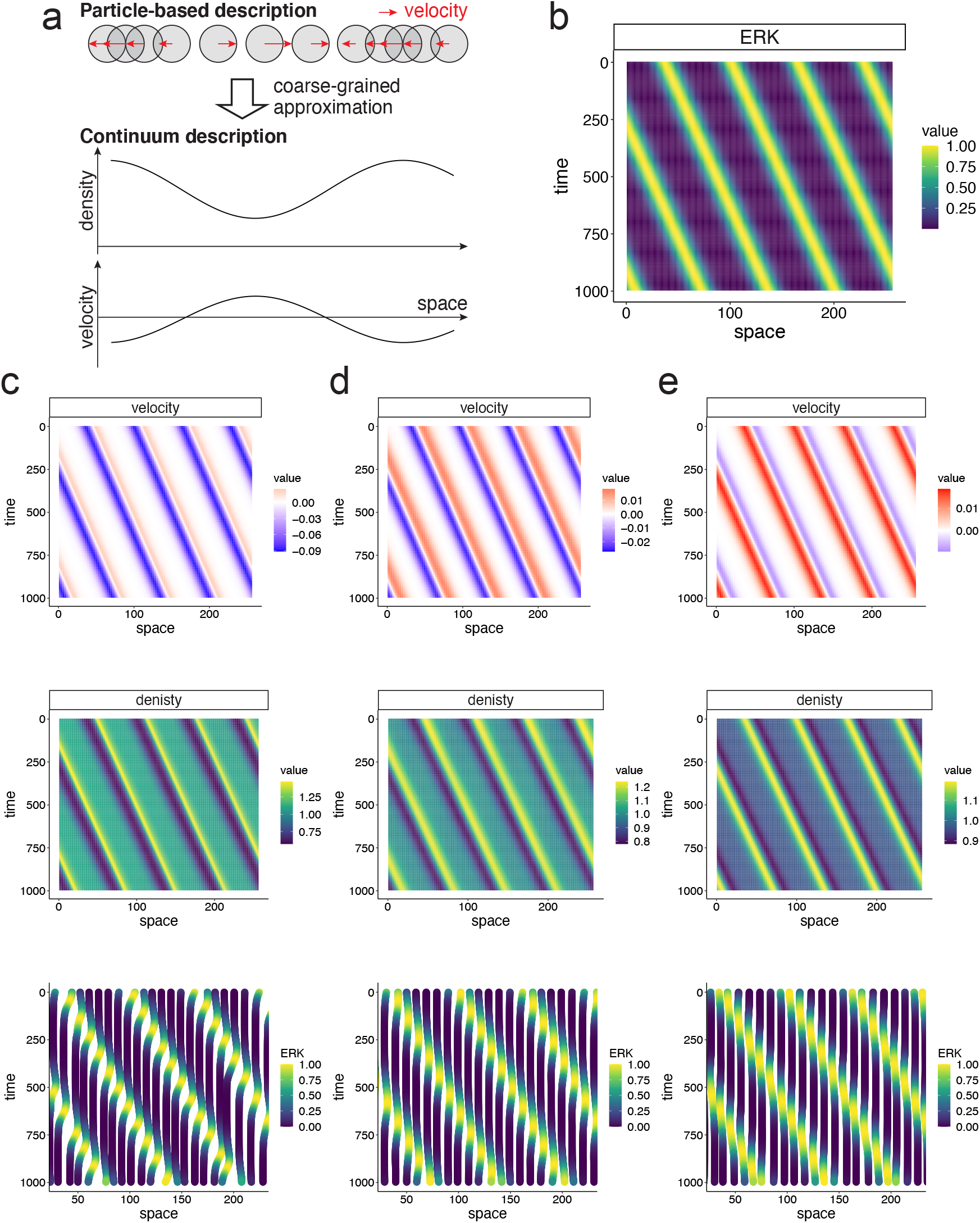
Simulations of the continuum model in 1D a. A schematic representation of the coarse-grained approximation from the particle-based model to continuum model. Migrations of particles, i.e., cells, are represented by density distribution and velocity. b. ERK traveling waves applied in simulations. c-e. Simulation results of velocity (upper panel), density (middle panel), and trajectories of the cells (lower panel). Parameters used in (c), (d) and (e) are the same as figure 2b, c and d, respectively.

In the coarse-grained description (6), collective cell migration is interpreted as a continuous flow field, expressed in Lagrangian description in fluid dynamics as follows:

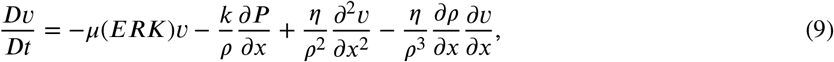

where *v* indicates velocity of the flow field, *P* = 1/*ρ*^∞^ − 1/*ρ* and *D*/*Dt* = *∂/∂t* + *v* grad (in this 1D case, *Dv/Dt* = *∂v/∂t* + *v∂v/∂x*, see Methods for details). Because *ρ*^∞^ = 1∕2*R*_0_(1 + *αERK*),

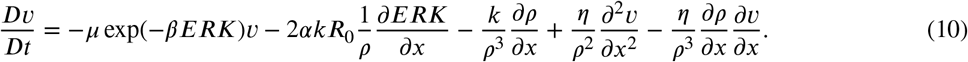

Note that *D*∕*Dt* is used for the Lagrangian description in which individual cells are tracked and the velocity change is detected as a function of time, whereas *∂*∕*∂t* is used for the Eulerian description in which velocity change is written as a function of space and time.

In contrast to equation (6), *v*, *P* and *ρ* are not indexed by *i*, since we addressed these variables as continuous functions in space and time. The cell density field also has a flux according to equation (8) and its dynamics are described by

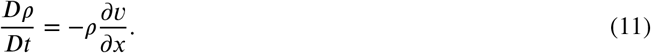

ERK waves were given the same function as those in the particle-based model (Fig 3b). With this continuum model, we performed simulations of fields of *v* and *ρ* in an Euler description with ERK waves that travel in a positive direction on the x-axis. The resulting velocity field showed negative values (Fig 3c), indicating flow in a direction opposite to that of the ERK waves. The migration in an opposite direction against ERK waves is consistent with our observations in previous experiments. As observed in the particle-based model, velocity fields showed the same direction values in some simulation parameters, as shown in (Fig 3c-e).

### Comparison between two models in 1D

To validate our continuum approximation in 1D, we compared the simulation results generated by the two models. To this end, we numerically reconstructed a trajectory of cellular migration, i.e., path line in fluid mechanics, from the simulated velocity field in the continuum model (Fig 3c-e) (see Methods). It should be noticed that the velocity field in the continuum model represents velocity at time *t* and space *x*, but does not correspond to tracking velocity of specific cells. By comparing the trajectories simulated in our two models, we confirmed that the trajectories generated by the two models were almost the same in various parameters (Fig 4). This indicates that our fluid approximation is valid and that the continuum model is able to explain ERK-dependent cell migration.

**Figure 4:**
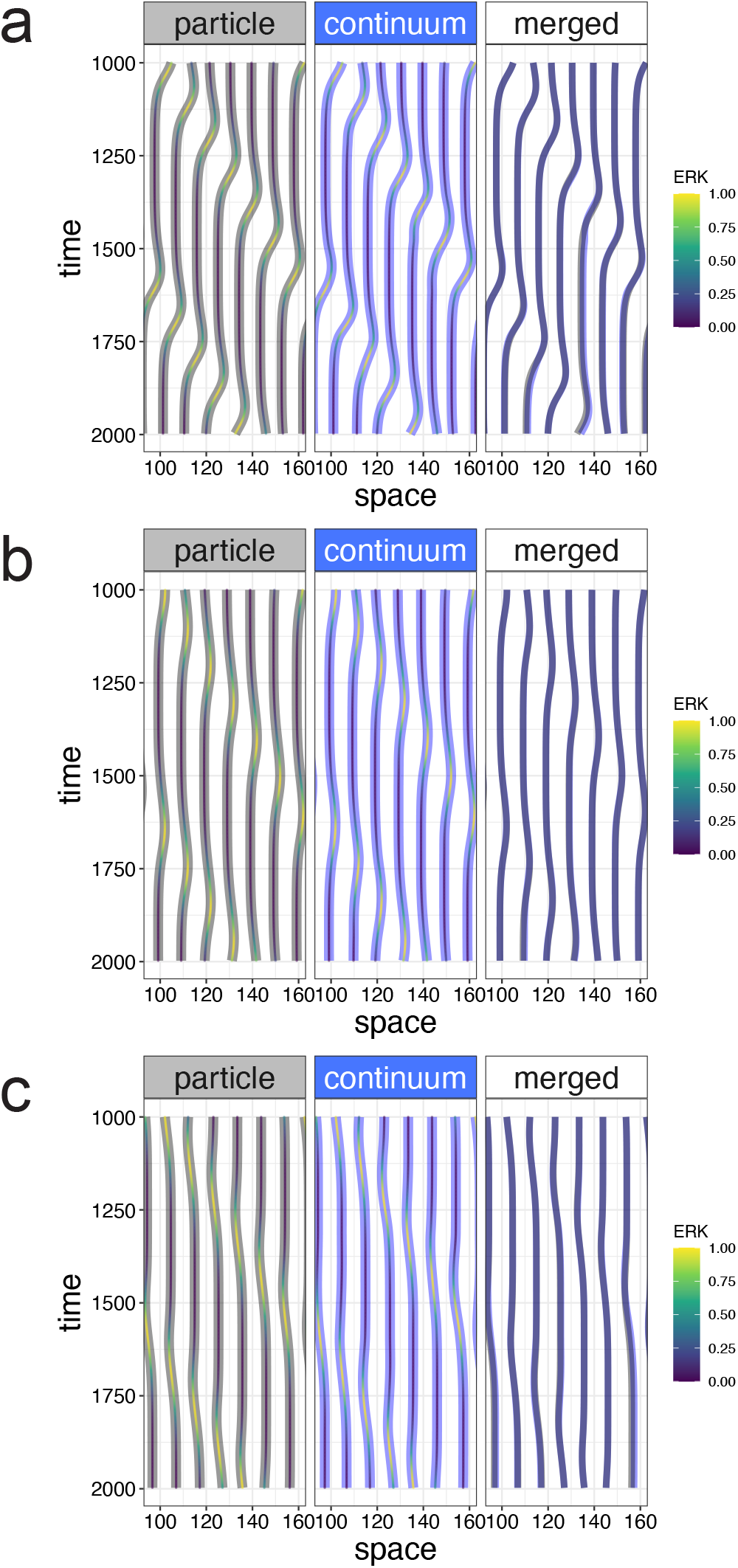
Comparison between the particle-based and continuum models in 1D simulation Comparisons between the migration trajectories simulated by two models with the same parameter values. In the left and center panels, the shaded gray and blue lines indicate trajectories simulated by the particle-based and continuum models, respectively, and the color of the inner lines indicates ERK activity. In the right panels, the shaded gray and blue lines were merged for comparison. Parameters used in (a), (b), and (c) are the same as figure 2b, c, and d, respectively.

### Comparison between two models in 2D

We further showed that our continuum model was valid not only in 1D but also in 2D. The 2D particle-based model was extended in 2D. The dynamics of a cell *i* at the position **x**_*i*_ = (*x*_*i*_, *y*_*i*_) are described by following ODEs:

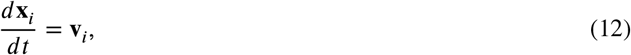

where **v**_*i*_ = (**v**_*x,i*_, **v**_*y,i*_) represents velocity of the cell *i*. Its dynamics are described as

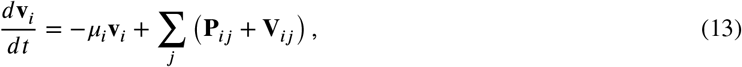

where *μ*_*i*_ indicates the friction coefficient, which is the same as that in equation (4), and **P**_*ij*_ and **V**_*ij*_ indicates pressure and viscosity, which a neighboring cell *j* applies to the cell *i*. They are expressed as

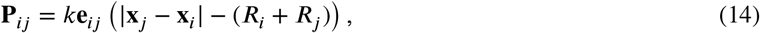

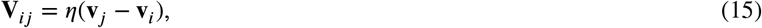

where *k*, *R*_*i*_, *μ*_*i*_ are the spring coefficient, natural length of a cell *i*, and friction coefficient, respectively, and **e**_*ij*_ is a unit vector (**x**_*j*_ − **x**_*i*_)∕|**x**_*j*_ − **x**_*i*_| from cell *i* to *j*. The neighboring cells were detected by calculating Voronoi diagrams.

In the 2D continuum model, our model was extended as follows:

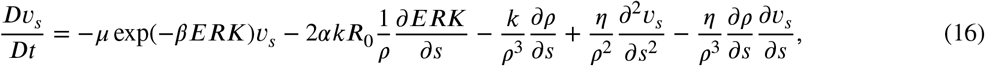

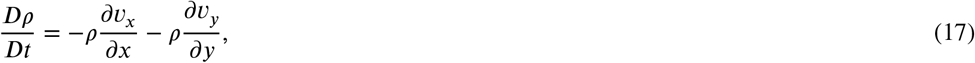

where *s* ∈ {*x, y*}. In both models, ERK was described as

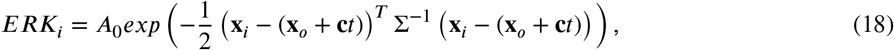

where Σ is a positive symmetric matrix regulating the shape of ERK waves.

With these equations, we conducted numerical simulations. In both models, we confirmed cellular migration in a ection opposite to that of traveling ERK waves (Fig 5a-c, supplemental movie 1-6). For comparison between the cellular trajectories simulated by two models in 2D, we plotted cellular trajectories in space-time coordinates, in which cellular trajectories within a 1D band region of interest were depicted (Fig 5d, e). We noticed that the trajectories in the particle-based model seemed less continuous than those in the continuum model, because the cells were in contact with discrete numbers of neighboring cells, e.g., five or six, and the neighboring pairs were frequently rearranged. Although the trajectories were disconnected because cells leave and enter the band region, the cells showed almost the same migration in a direction opposite to that of the ERK waves in both models. This result showed a clear consistency between the particle-based and continuum models, even in 2D. We further confirmed a consistency between the two models by comparing the simulated trajectories in 2D space (Fig 5d, e). Taken together, our continuum approximation was validated to recapitulate tissue dynamics, even in 2D.

**Figure 5:**
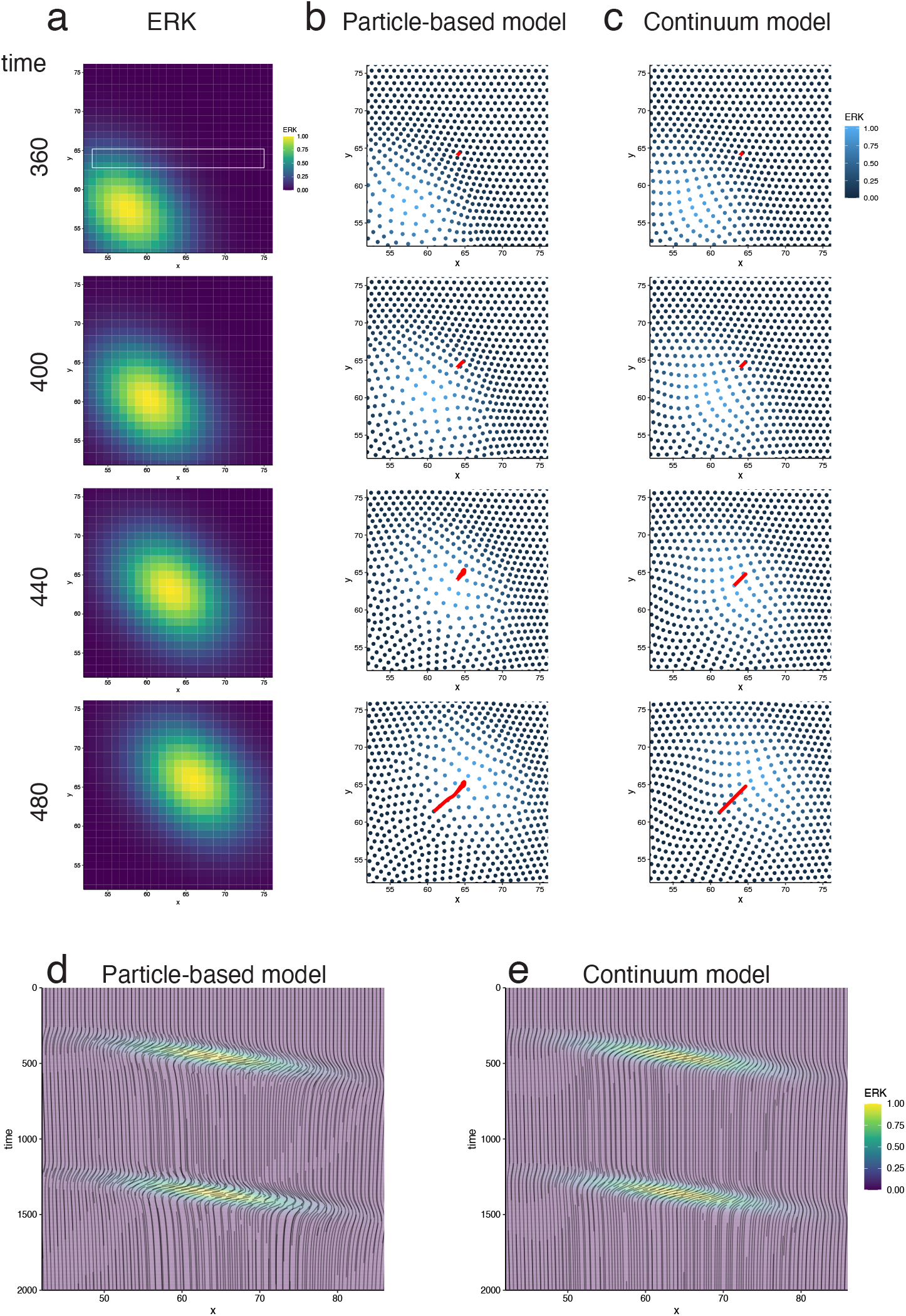
Comparison between the particle-based and continuum models in 2D simulation a. ERK wave applied in the simulations. b-c. Simulation results in the particle-based and continuum models. Each dot represents cells in the particle-based model (b) and virtual marker simulated in the Lagrange description (c). Red lines indicate the trajectories of the cell (b) and virtual marker (c), which are initially located at the same position. See a supplemental movie for these simulations. d-e. Trajectories of the simulated cells in space (*x*) - time coordinates in the particle-based (d) and continuum (e) models. Cellular migration trajectories within a white box region shown in a.

## Discussion

Here we proposed a hierarchical modeling approach to understand mechano-chemical epithelial dynamics, which acquires causality beyond the hierarchy between the cellular and tissue levels. We showed that the previous particle-based model at the cellular level can be hierarchically transformed into the continuum model to reveal tissue-level epithelial dynamics. Through numerical simulation, we demonstrated the clear consistency of the trajectories simulated by the continuum model and our previous particle model, and our continuum model successfully recapitulated our previous experimental observations. These results indicated the validity of our hierarchical modeling. Thus, our continuum model offers a new theoretical platform to reveal hierarchical regulatory systems between cells and tissues, integrating mechano-chemical signals that determine both cellular and tissue behaviors.

Several coarse-grained models have been developed [11–15], however, no models for the incorporation of mechanical dynamics with molecular activities that were quantitatively measured by recent imaging techniques with the FRET-based biosensor have been introduced. Due to this lack of integration, mechanistic insights into how intracellular signals, e.g., MAPK and WNT, regulate the mechanical dynamics of tissues and cells have been limited [16], in spite of the recent progress in imaging techniques. Therefore, our model is the first coarse-grained model to integrate the complex regulatory mechanisms of mechano-chemical tissue dynamics.

Although the epithelial tissue is a 2D sheet, our previous particle-based model was restricted to 1D observations. the other hand, our continuum model can easily be extended to function as a two-dimensional model and reflects situ 2D epithelial sheet dynamics observed by imaging experiments. Furthermore, because the continuum model coarse-grains noisy rearrangement of cells, our hierarchical modeling approach presents a new theoretical methodology to understand tissue dynamics without monitoring all cellular behaviors. Therefore, the continuum model is more advantageous than other models at the cellular level, in terms of enabling the interpretation of 2D epithelial sheet dynamics observed by the imaging data [17–19].

Since the recent progress in live cell imaging techniques has led to an increase in the number of signaling molecules hat can be monitored at the same time, future works need to elucidate the importance of the chemical factors other han ERK. Our continuum model currently takes only one chemical factor into account, and it needs to be further extended to consider two or more factors. Along with the multiple-chemical signal quantifications, recent progress in the quantitative biology field has enabled the measurement of mechanical signals such as pressure and density [20, 21]. No methods had been introduced, however, to analyze mechanical signals with canonical molecular biochemical signals at the same time. Our continuum model provides a new theoretical basis to analyze the cellular system, integrating both mechanical and chemical signals.

## Methods

### Detailed derivation of the continuum model

Equation (2) of the particle-based model is rewritten as

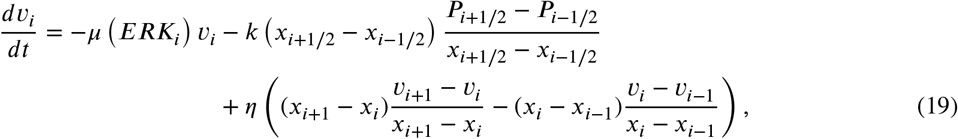

where *P*_*j*_ = (*R*_*j*+1∕2_ + *R*_*j*−1∕2_) − (*x*_*j*+1∕2_ − *x*_*j*−1∕2_) and *P*_*i*+1∕2_ and *P*_*i*−1∕2_ correspond to pressure from neighboring cells on the right and left, respectively. Here, we defined the cell density at cell *i* as *ρ*_*i*_ = 1∕(*x*_*i*+1∕2_ − *x*_*i*−1∕2_), where *i* + 1∕2 represents the middle point index between cell *i* and cell *i* + 1. In the same way, the natural cell density at cell *i* is defined by the inverse of the sum of the natural radius of neighboring cells as 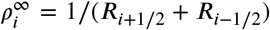. These definitions lead to 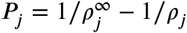 Thus, the equation (2) becomes

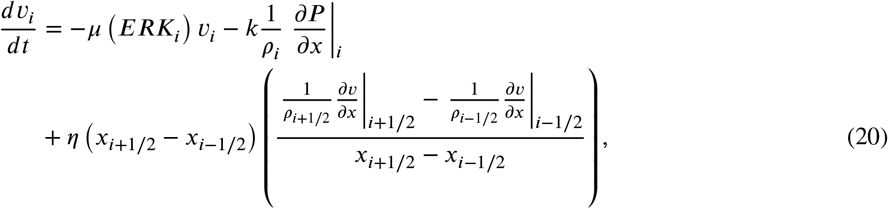

where

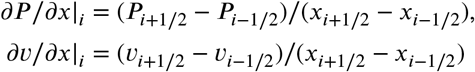

According to continuous approximation, i.e. 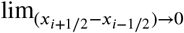, this equation is transformed into

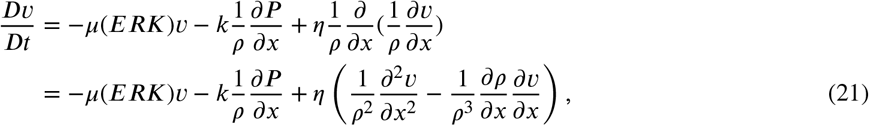

where *v* indicates continuous function of *x* and *t* as *v*(*x, t*), *P* = 1∕*ρ*^∞^(*ERK*) − 1∕*ρ*, *ρ*^∞^(*ERK*) = 1∕2*R*(*ERK*) and *Dv*∕*Dt* = *∂v*∕*∂t* + *v ∂v*∕*∂x*. Note that *Dv*∕*Dt* indicates Lagrange description of velocity, representing the temporal ivative the mig ation elocity of tr king cells. *P* is defined by *P*(*x, t*) = 1∕*ρ*^∞^(*x, t*)−1∕*ρ*(*x, t*), where *ρ*^∞^(*x, t*) = 1∕2*R* (*ERK*(*x, t*)). Here, we considered *R*(*ERK*) = *R*_0_(1 + *αERK*) in the particle-based model, which leads to equation (10).

To obtain the continuum dynamics of cell density, we consider the temporal derivative of *ρ*_*i*_ referring to the definition *ρ*_*i*_ = 1∕(*x*_*i*+1∕2_ − *x*_*i*−1∕2_) as

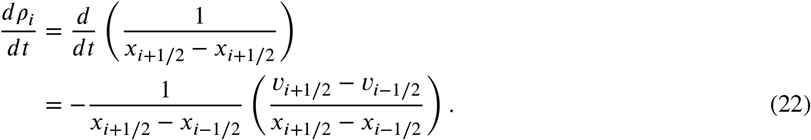

As described for velocity above, this equation changes to

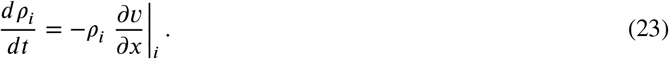

Again, continuum approximation leads to temporal derivative of density in the Lagrange description:

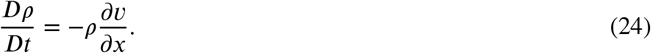

This equation can be changed in Euler description as

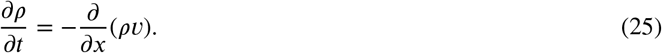

This equation is well known as the continuity equation, resulting from the law of mass conservation in continuum dynamics. Because it is naturally derived, our continuum approximation was accurate.

### Numerical simulation for the particle-based model

Numerical simulations for the particle-based models in 1D and 2D were conducted by the Runge-Kutta method with fourth order accuracy. In the 1D simulation, the initial positions of the particles were equally spaced with unit intervals. In the 2D simulation, the initial positions of the particles were equally spaced on the xy-plane in a hexagonal lattice with unit intervals. Neighboring cells were detected according to the Voronoi diagram with a C++ package qhull [22].

### Numerical simulation for the continuum model

#### Discretization of the continuum model

For numerical simulations, the continuum PDE model in Euler description was discretized with local Lax-Friedrichs flux [23]. PDEs were generally expressed by

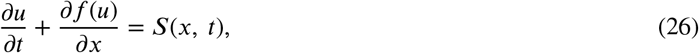

where *u* ∈ {*ρ*(*x, t*), *v*(*x t*)}, *f*(*u*) indicates flux, e.g., *f*(*ρ*) = *ρv* for cell density and *f*(*v*) = *v*^2^∕2 for cell velocity, and *S*(*t*) indicates reaction terms, e.g., *S*(*t*) = 0 for cell density and *S*(*t*) equals to the right-hand-side of equation (10).

This PDE can be discretized in space and time as

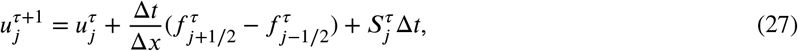

where *j* and *τ* denote indices of discretized space and time, respectively; *j* ± 1∕2 corresponds to mid positions between *j* and *j* ± 1; Δ*x* and Δ*τ* indicate the discretized intervals of space and time, respectively; 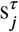 indicates value at time *τ*Δ*t* and position *j*Δ*x*, (*s* ∈ {*u, f, g*}); 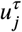 represents the spatial average of *u*(*x*) at space *j* as

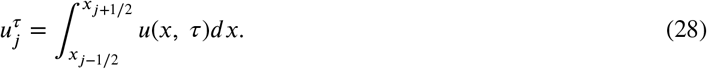

Since *f*_*j*±1∕2_ indicates flux at the mid positions between *j* and *j* ± 1, this equation satisfies the conservation of *u*. Here, we considered the discontinuity at *j* ± 1∕2 such that

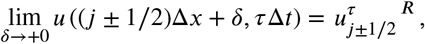

and

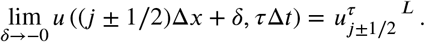

Fluxes at *j* ± 1∕2 were defined by Lax-Friedrichs flux as

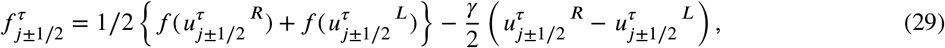

where *γ* = max_*x*_|*f*(*u*(*x*))|. The first term represents the average flux, whereas the second term indicates artificial diffusion for avoiding numerical stability. 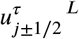 and 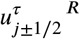 in equation (29) were calculated by weighted essentially non-oscillatory (WENO) scheme [23, 24] described below.

#### The WENO scheme

We used the fifth-order WENO scheme to compute 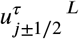 and 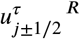 from values of *u* at the grid points in the stencils *S*^*L*^ = {*x*_*j*−2_, *x*_*j*−1_, *x*_*j*_, *x*_*j*+1_, *x*_*j*+2_} and *S*^*R*^ = {*x*_*j*−1_, *x*_*j*_, *x*_*j*+1_, *x*_*j*+2_, *x*_*j*+3_}, respectively. The WENO scheme is a local interpolation method that takes into account the discontinuity of *u*(*x*). Because *u_j_* represents the spatial average of *u*(*x*) over the interval [*x*_*j*−1∕2_, *x*_*j*+1∕2_], we just know its integral as

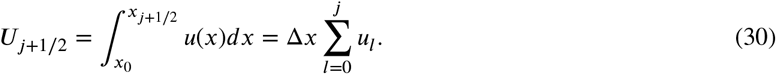

In the WENO reconstruction procedure [24], values of U were interpolated by the three different stencils of

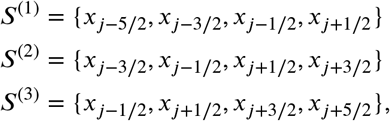

as third-order polynomials, *P*^(1)^(*x*), *P*^(2)^(*x*) and *P*^(3)^(*x*), respectively. Then, values of *u* were reconstructed by second-order polynomials as *p*^(*n*)^(*x*) = *dP*^(*n*)^(*x*)∕*dx*, and *u*_*j*+1∕2_ is approximated to values of *p*^(*n*)^(*x*_*j*+1∕2_) as

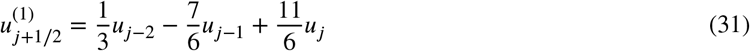

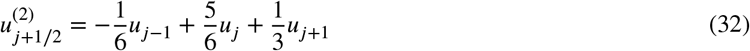

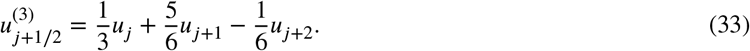

Then, *u*_*j*+1∕2_^*L*^ was approximated by a convex combination of the three reconstructed *u*_*j*+1∕2_ as

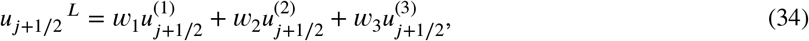

where ∑_*n*_ w_*n*_ = 1 and these weights are modulated in a discontinuity-dependent manner as

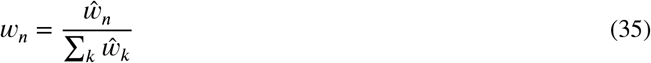

with

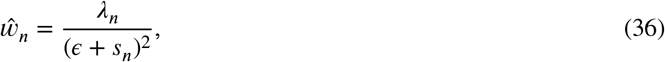

*λ*_1_ = 1∕10*, λ*_2_ = 6∕10*, λ*_3_ = 3∕10 and *є* = 10^−6^. *s*_*n*_ is smoothness indicator of *p*^(*n*)^ (*x*) defined by

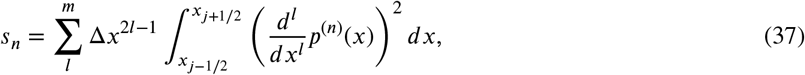

where *m* is the polynomial degree of *p*(*n*)(*x*) (in this case, *m* = 2). The smoothness indicators were formulated by

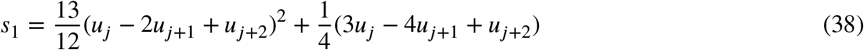

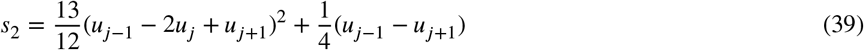

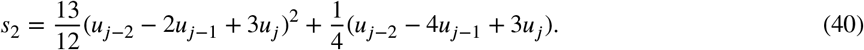

Note that if *u*(*x*) is smooth from *x*_*j*−1_ to *x*_*j*+3_ (*i.e., s*_1_ = *s*_2_ = *s*_3_), *u*_*j*+1∕2_^*L*^ becomes exactly the same as a polynomial of degree at most four on large stencil *S*. Similarly, *u*_*j*+1∕2_^*R*^can be calculated on stencil *S*^*R*^, but in a mirror symmetric manner with respect to *x*_*j*_ of the above procedure.

#### Numerical simulation in Euler description

The equation (26) can be written as a method-of-line ODEs system

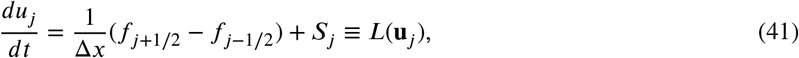

where **u**_*j*_ = {*u*_*j*−2_, *u*_*j*−1_, *u*_*j*_, *u*_*j*+1_, *u*_*j*+2_}. These ODEs were temporally integrated by the total variation diminishing (TVD) Runge-Kutta method with third order accuracy [23] as

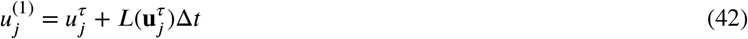

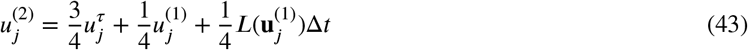

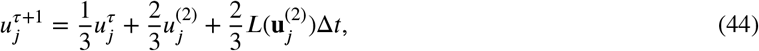

where 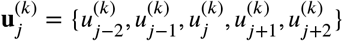.

#### 2D numerical simulation in Euler description

For a 2D simulation in Euler description in the WENO scheme [23], PDEs were generally expressed by

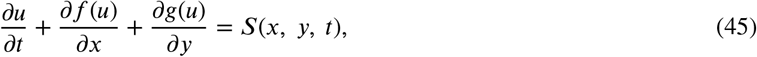

where *S*(*x, y, t*) expresses the reaction term. In the 2D case, we calculated momentum *ρv*_*x*_ and *ρv*_*y*_ as *u* in the equation (45) instead of calculating *v*_*x*_ and *v*_*y*_ directly. In *x* direction, *u*, *f*(*u*), *g*(*u*) and *S*(*x, y, t*) in the equation (45) were

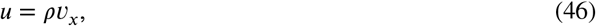

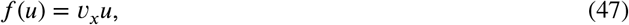

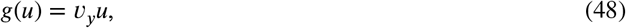

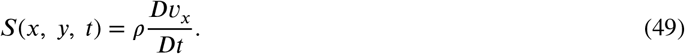

With equations (46) to (48), each term on the left-hand-side in equation (45) becomes

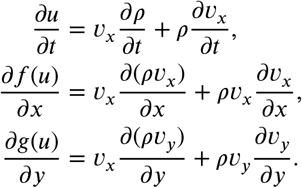

The sum of the first terms on the right-hand-side of the equation was removed as,

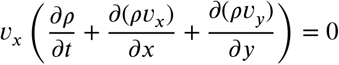

since the equation of continuity appears in the brackets. The sum of the second terms becomes

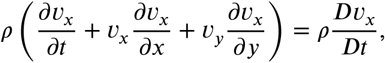

which corresponds to the equation (49).

Then, the equation (45) was discretized in a finite different scheme [23] as

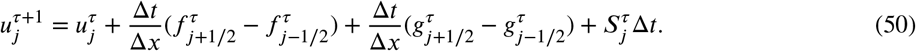

Note that in the finite different scheme, each of the numerical fluxes 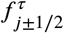 and 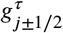 can be computed independently.

Therefore, the numerical fluxes were computed in the same way as in the 1D case.

#### Numerical simulation in Lagrange description

After obtaining the field value *v* (*x, t*), Lagrange-described value *x*(*x*_0_*, T*) at time T is calculated by

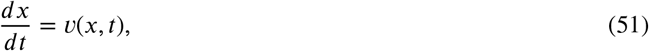

by referring *v* (*x, t*). The calculations were conducted using the Runge-Kutta method with fourth order accuracy in 1D simulations, and on an implicit Runge-Kutta method of the Radau IIA family of order five in 2D simulations. Initial positions of particles on the tissue were determined as the same initial positions as those in the particle model simulations.

## Data availability

All relevant data have been provided within the paper.

## Code availability

The codes will be available in GitHub when the manuscript would be accepted.

## Acknowledgement

We thank Dr. Michiyuki Matsuda from Kyoto University for his helpful discussions with us about the paper and Dr. Yangjin Kim from Konkuk University for his suggestion to use the WENO scheme. This study was mainly supported by the Cooperative Study Program of Exploratory Research Center on Life and Living Systems (ExCELLS) (ExCELLS program No.18-201). It was also supported by JSPS KAKENHI Grants (16K16147 (H.N.), 18H02444, 19H05798 (K.A.), 19H05438, 19K16207 (Y.K.), 20J15811 (Y.A.)), by JST CREST Grant JPMJCR1654 (K.A.).

## Author contributions

H.N. conceived the project. Y.A., H.N. and Y.K. developed the model. Y.A. implemented and performed numerical simulations. Y.A. and K.A. analyzed data. Y.A. and H.N. wrote the manuscript with input from all authors.

---

**Supplemental figure 1:**
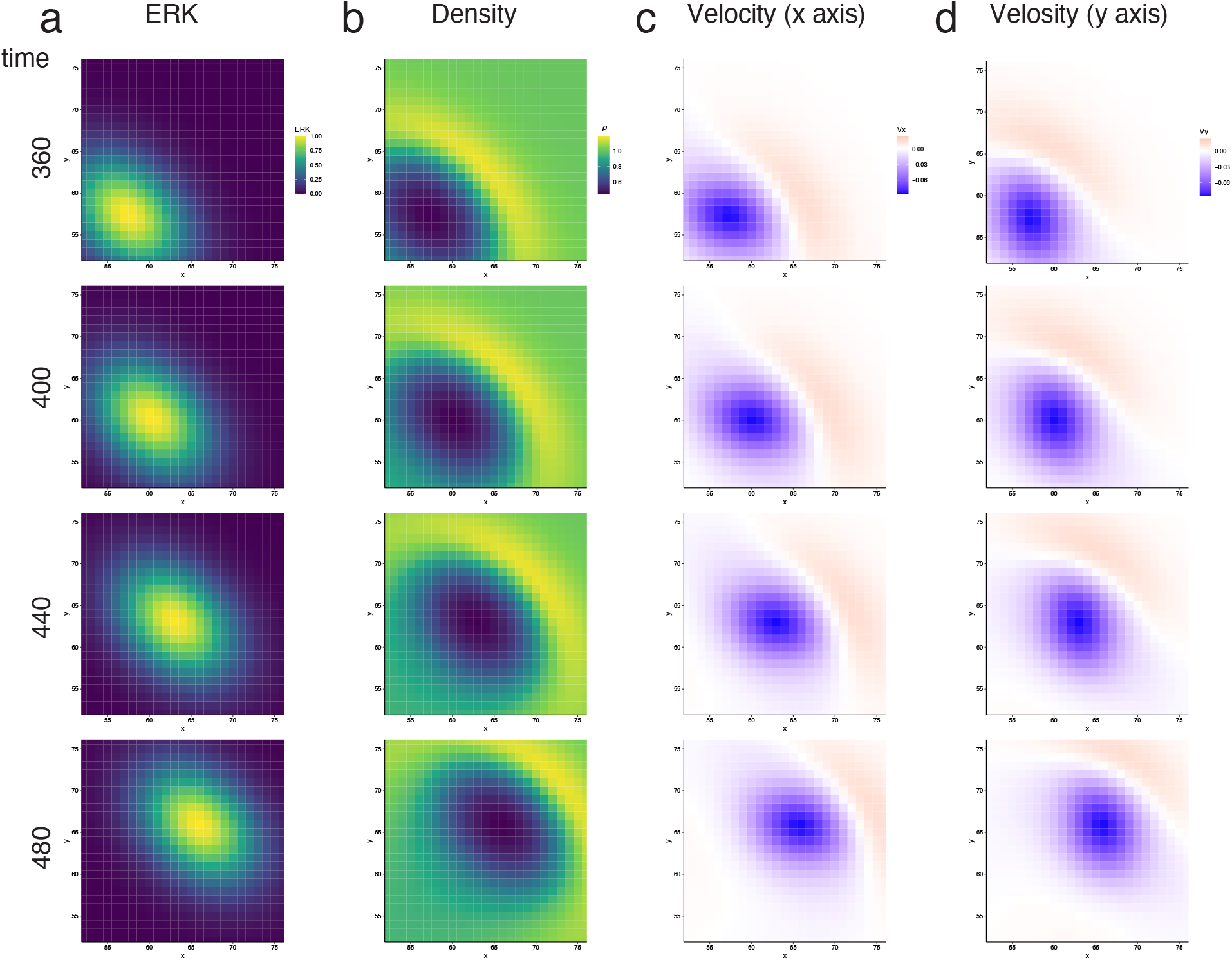
Two-dimensional simulation in the continuum model a-d. Simulated results in Euler description. ERK wave (a), Cell density (b), Velocities on x-direction (c) and y-direction (d). See a supplemental movie for the simulation.

